# Regulator of G Protein Signaling 6 Negatively Regulates Platelet Activation and Arterial Thrombosis in Mice

**DOI:** 10.64898/2026.04.23.720481

**Authors:** Renat Roytenberg, Boyd R. Rorabaugh, Hong Yue, Robbie Jividen, Scott J. Cameron, Wei Li

## Abstract

**Background:** Platelet activation via G protein-coupled receptors (GPCRs) is central to arterial thrombosis. P2Y12 is a canonical Gi-coupled receptor mediating ADP-dependent platelet activation, yet the role of Regulator of G protein Signaling 6 (RGS6), a modulator of Gi signaling, in platelet function and thrombosis remains unclear.

**Objectives:** To determine the role of RGS6 in platelet activation and arterial thrombosis and to define its impact on P2Y_12_/Gi signaling.

**Methods:** Arterial thrombosis was assessed using a FeCl□-induced carotid artery injury model in wild-type (WT) and *Rgs6*^*–/–*^ mice. Platelet aggregation was measured ex vivo. Signaling pathways were analyzed by Western blot in ADP-stimulated platelets. P2Y_12_/Gi signaling was further evaluated using a cAMP-responsive luciferase reporter assay in HEK293 cells.

**Results:** Male *Rgs6*^*–/–*^ mice exhibited significantly accelerated thrombosis compared with WT controls. *Rgs6*^*–/–*^ platelets showed enhanced ADP-induced aggregation, whereas collagen-induced aggregation was unchanged. In ADP-stimulated platelets, RGS6 deficiency altered signaling kinetics, characterized by delayed Akt phosphorylation and reduced PKA and VASP phosphorylation. In a heterologous cAMP-luciferase assay, RGS6 attenuated P2Y_12_/Gi-mediated suppression of cAMP. Two-way ANOVA demonstrated significant effects of ADP and RGS6 expression on luciferase activity, with no interaction, indicating that RGS6 modulates signaling magnitude rather than agonist sensitivity. Pharmacologic inhibition of P2Y_12_ with clopidogrel abolished the genotype-dependent difference in thrombosis in vivo.

**Conclusions:** RGS6 acts as a negative regulator of platelet P2Y_12_/Gi signaling and thrombus formation. Loss of RGS6 enhances ADP-dependent platelet activation and accelerates arterial thrombosis, establishing RGS6 as an endogenous brake on platelet activation.

## Introduction

Thrombotic events are a major cause of morbidity and mortality in the US and worldwide [1-3]. Platelet activation and increased reactivity at the site of vascular injury are the primary pathogenic components of thrombosis, which lead to vessel occlusion and causing myocardial infarction and ischemic stroke. Platelets have many surface receptors and, when activated, promoting platelet adhesion and initial activation in response to exposed collagen at the site of vascular injury [4-6]. Adherent, activated platelets release soluble agonists (e.g., ADP, thrombin, and thromboxane A_2_ [TXA2]), which locally activate additional platelets via surface G protein-coupled receptors (GPCRs). The major platelet GPCRs that mediate platelet activation include ADP receptors P2Y_12_ and P2Y_1_, thrombin receptors protease-activated receptor (PAR) 1 and PAR4, and TXA2 receptor (TP) [5, 7, 8]. Consequently, the FDA has approved aspirin (non-selective COX inhibitor, which inhibits TXA2 production and thus reduces TP agonism), clopidogrel and prasugrel (P2Y_12_ inhibitors), and vorapaxar (PAR1 inhibitor) as anti-platelet/thrombotic drugs, which are clinically indicated for both primary or secondary prevention of platelet-mediated thrombotic events [9-12]. However, these drugs have major side effects, including injury to the gastrointestinal mucosa, thrombocytopenia, and systemic hemorrhage [3, 9-12]. In addition, some patients do not respond to these regimens and have a high incidence of recurrent thrombosis [3]. Therefore, defining the molecular mechanisms for platelet activation in thrombus formation remains a significant clinical unmet need.

ADP and thrombin receptors are coupled to three major G proteins: G_αq_, G_α12/13_, and G_αi_ [13, 14]. ADP-induced platelet activation requires concomitant signaling from both P2Y_1_ and P2Y_12_ receptors that couple to G_αq_ and G_αi_, respectively. Regulators of G protein signaling (RGS) proteins enhance the rate of GTP hydrolysis by the Gα subunit of heterotrimeric G proteins. This provides a mechanism for cells to control the magnitude and duration of signaling through GPCRs [15, 16]. Twenty canonical RGS proteins have been identified [17]. Members of the RGS protein family exhibit unique tissue distributions and selectivity for different Gα isoforms, making them attractive therapeutic drug targets. Previous work has demonstrated that RGS10, RGS16, and RGS18 are expressed in platelets where they regulate platelet activation and thrombosis [18-22]. RGS6 has also been identified in platelets at both the mRNA [23, 24] and protein levels [25, 26]. Notably, platelet RGS6 expression was reduced by approximately 30% in patients with COVID-19, irrespective of ICU admission status [23]. COVID-19 patients are a high-risk cohort to develop thrombosis. Interestingly, a recent pharmacogenomic study in Caribbean Hispanic patients identified an intronic RGS6 variant that is associated with clopidogrel responsiveness, further supporting a role for RGS6 in platelet P2Y_12_ signaling [27]. However, the function of RGS6 in platelets has not been explored. In this study, for the first time, we demonstrated that RGS6 regulates ADP-induced platelet aggregation and is an important regulator of thrombosis following vascular injury.

## Methods

### Animals

*Rgs6*^*–/–*^ mice were a generous gift from Dr. Rory Fisher (University of Iowa School of Medicine, Iowa City, Iowa). The generation of these mice has been previously described [28]. The *Rgs6*^*+/–*^ mouse strain has been backcrossed onto the wildtype (WT) C57BL6/J (Jackson Labs, Bar Harbor, Maine; stock #000664) background more than 15 generations. *Rgs6*^*–/–*^ mice were maintained in homozygous inbreeding. WT (*Rgs6*^*+/+*^) mice purchased from The Jackson Labs were bred and housed within the same room as the *Rgs6*^*–/–*^ mice and used as controls. All procedures and manipulations of animals have been approved by the Institutional Animal Care and Use Committee of Marshall University (#: 1033528, PI: WL, and #: 1174948, PI: BR).

### Materials

Platelet agonists including ADP (P/N 384) and collagen (P/N 385) were purchased from Chrono-log (Havertown, PA). Antibodies to phosphorylated AKT (4060S), pan-AKT (2920S), phosphorylated PKA (4781S), pan-PKA (5842S), HRP-conjugated Pan-actin (12748S), HRP Conjugated anti-Rabbit (7074S) or mouse (7076S) IgG secondary antibody were purchased from Cell Signaling Technology (Danvers, MA). Antibodies to P2Y_12_ (ab184411, ab183066) were purchased from abcam (Waltham, MA). All other chemical reagents were purchased from Millipore Sigma (Burlington, MA) except where specifically indicated.

### Murine FeCl_3_-injury-induced carotid artery thrombosis model

The ferric chloride (FeCl_3_)-injury-induced carotid artery thrombosis model has been described previously [29, 30]. Briefly, mice of both sexes, 8 to 16 weeks old, were anesthetized by a mixture of ketamine/xylazine (100/10 mg/kg) via intraperitoneal injection. Platelets were labeled through direct jugular vein injection of 100 μl of rhodamine 6G solution (Sigma 252433-1G, 0.5 mg/ml in saline, 0.2 μm filtered). The carotid artery was exposed, and injury was induced by topically applying a piece of filter paper (1 x 2 mm) saturated with 7.5% FeCl_3_ solution for 1 minute. Thrombus formation was observed in real-time using intravital microscopy with a Leica DM6 FS fluorescent microscope (Deerfield, IL, USA) attached to a Gibraltar Platform. Video imaging was conducted using a QImaging Retiga R1: 1.4 Megapixel Color CCD camera system with mono color mode (Teledyne Photometrics, Tucson, AZ, USA) and StreamPix version 7.1 software (Norpix, Montreal, Canada). The endpoints were set as 1) blood flow has ceased for > 30 seconds, or 2) occlusion is not seen 30 minutes after FeCl_3_ injury. In the second case, 30 minutes was assigned as the data value for statistical analysis.

A different cohort of mice were fed a clopidogrel solution (clopidogrel bisulfate in ethanol, serially diluted in saline) via oral gavage at 1 mg/kg (uncertain dose to produce a significant antithrombotic effect) [31], once per day for 72h before experimentation. The operators were blinded to animal genotypes.

### Tail bleeding assay

Tail bleeding assay was conducted as previously described [31, 32]. Briefly, 10-12-week-old male mice were anesthetized with ketamine/xylazine (100/10 mg/kg). One centimeter from the tail tip was transected using a sharp scalpel, and the tail was immediately immersed into 37 °C warm saline. Bleeding time was recorded from the moment of tail transection until bleeding completely ceased, which is named as 1^st^ bleed. After the 1^st^ bleeding, the wound was nicked with the scalpel, the tail was immediately immersed in a new tube containing warm saline, and the time to bleeding cessation was recorded, which is designated as the 2^nd^ bleed.

If bleeding persisted for longer than 6 minutes (3 times the mean bleeding time in WT mice), a gentle pressure was applied to the tail for 2 minutes to achieve hemostasis. For statistical analysis, a bleeding time of 6 minutes was assigned to these mice.

### Whole blood count and platelet isolation

Mice were anesthetized with ketamine/xylazine (100/10 mg/kg), and 0.9 - 1 mL whole blood was collected through inferior vena cava puncture using 0.109 M sodium citrate as an anticoagulant. Blood cells were counted immediately using Hemavet 950FS. Modified Tyrode’s buffer (concentration of components in mM: 137 NaCl, 2.7 KCl, 12 NaHCO_3_, 0.4 NaH2PO_4_, 5 HEPES, 0.1% glucose, and 0.35% BSA, pH 7.2) was added to the collected whole blood at 0.7 volumes, mixed, and then platelet-rich plasma (PRP) was isolated by centrifugation at 100 g for 10 min. The sediment containing red blood cells and leukocytes was further centrifuged at 13,000 rpm for 1 min to isolate platelet-poor plasma (PPP) [32].

### Platelet aggregation assay

The platelet concentration in PRP was counted with a hemocytometer and adjusted to 2.5E+08/mL with PPP, and 0.4 ml of this platelet suspension was used for the platelet aggregation assay using Chrono-log 700 with an agitation speed of 1,200 rpm. CaCl_2_/MgCl_2_ was repleted at a final concentration of 1 mM immediately before adding a platelet agonist [31-34].

### Evaluation of P2Y_12_-mediated platelet signaling activation

Platelets in PRP (2.5E+08/ml) were pooled from 6 mice, divided into 4 aliquots, and then stimulated with 2.5 μM ADP for 0, 1, 3, or 5 min [34]. Platelet activation was stopped by adding a final concentration of 1 mM EDTA and 0.5 µM PGEL to the reaction mixture. The platelets were then pelleted by centrifugation at 13,000 RPM for 15 seconds, immediately lysed in radioimmunoprecipitation assay (RIPA) buffer containing 1 x Halt™ Protease and Phosphatase Inhibitor Single-Use Cocktail, EDTA-Free (100 X stock solution, ThermoFisher, Cat# 78443). Protein concentration was determined using a Bio-Rad Protein assay. AKT, PKA, and VASP phosphorylation were assessed by immunoblotting assays.

### cAMP-luciferase reporter assay

The GloResponse™ CRE-luc2P HEK293 cell line was purchased from Promega (Cat# 8500, Madison, WI) and has been used in previous studies [35]. HEK293 cells were seeded at 10^5^ cells/well in 24-well plates in Dulbecco’s Modified Eagle Medium (DMEM) supplemented with 10% fetal bovine serum (Atlanta Biologicals Inc, Flowery Branch, GA, USA) and 1 X Antibiotic-Antimycotic (Gibco). Six hours after seeding, cells were transfected using Lipofectamine™ 3000 (Cat# L3000-015, Thermo Fisher Scientific, USA) according to the manufacturer’s protocol. Cells were transfected with pcDNA3.1-P2Y_12_, pcDNA3.1-*Rgs6* (cDNA Resource Center, Bloomsburg, PA), or their combination, along with Renilla plasmids. Empty pcDNA3.1 vector was used to equalize the total DNA amount (500 ng/well) transfected.

After 24h, the medium was replaced with serum-free DMEM. Following a 4h incubation in serum-free DMEM, cells were treated with varying concentrations of ADP or 0.9% saline for 1h at 37°C. Cells were then lysed in passive lysis buffer provided by the Dual-Luciferase® Reporter Assay System (Cat #: E1910, Promega, USA). Bioluminescence readings for firefly luciferase and Renilla were obtained using a SpectraMax L microplate reader operated with SoftMax Pro7 software (Molecular Devices). Luciferase activity was normalized to the corresponding Renilla signal and further normalized with the average value of cells transfected with empty pCDNA3.1.

### Statistics

Data are expressed as mean ± SEM. Results were analyzed by 2-tailed Student’s *t* test, Mann Whitney test, or 2-way/1-way ANOVA with Bonferroni post-hoc test for multiple comparisons using GraphPad Prism (version 10.2.2). In some cases, data were analyzed by Log-rank test using the Kaplan-Meier survival curve. *P* < 0.05 was considered statistically significant.

## Results

### *Rgs6* gene deficiency significantly enhanced thrombosis in male mice

To evaluate whether RGS6 regulates thrombosis, we employed the 7.5% FeCl□-induced carotid artery injury model [29, 30] to assess in vivo thrombus formation in WT and *Rgs6*^*–/–*^ mice. As shown in **Fig.1A**, loss of RGS6 did not appreciably alter the initial phase of platelet adhesion and aggregation at the site of vascular injury, as observed 1 minute after injury. In contrast, the subsequent wave of platelet accumulation was markedly accelerated in *Rgs6*^*–/–*^ male mice, resulting in a significantly shorter time to occlusive thrombus formation compared with WT controls. Overall, RGS6 deficiency accelerated the time to form an occlusive thrombus in male mice (**Fig. 1B**). Interestingly, while it was not statistically different, *Rgs6* deficiency in females tended to prolong the time to form an occlusive thrombus (**Fig. 1C**). However, due to the absence of statistical significance in females, all subsequent experiments were conducted using only male mice.

**Fig. 1.**
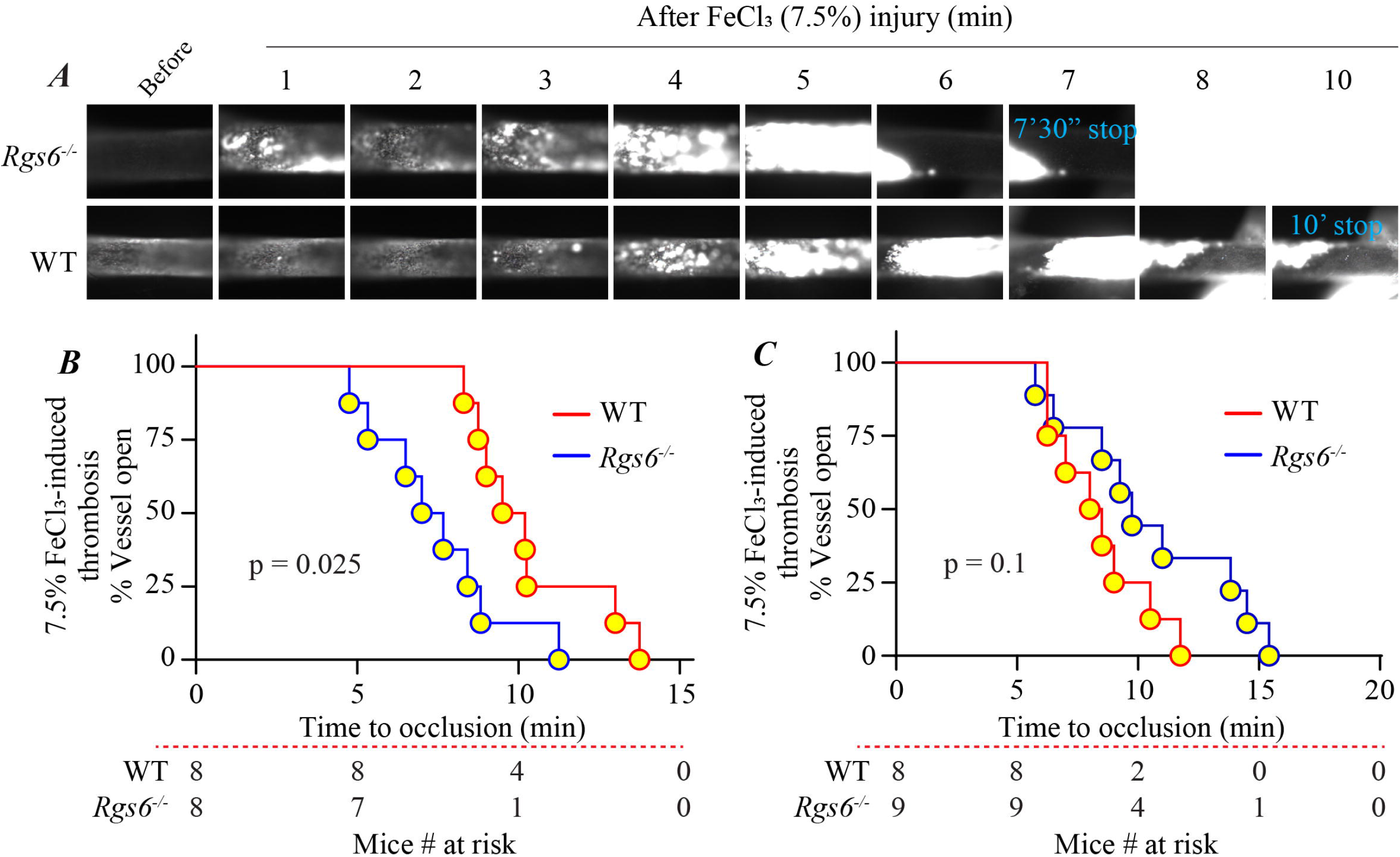
RGS6 deficiency enhances arterial thrombosis in male mice, but not in females. (**A**) Representative video images of thrombus formation in the carotid artery of male mice following 7.5% FeCl_3_ treatment. Platelets were labeled via direct intravenous injection of Rhodamine 6G. (**B**) Kaplan-Meier curve showing the time to occlusive thrombus formation after injury in male mice. (**C**) Kaplan-Meier survival curve data showing the time to occlusive thrombus formation after injury in female mice.

RGS6 deficiency did not affect complete blood cell counts (**Table 1**), suggesting that the prothrombotic phenotype observed in male mice was not attributable to alterations in circulating blood cell components. RGS6 deficiency also did not affect tail bleeding time (**Supplemental Fig. I**).

**Table 1.**
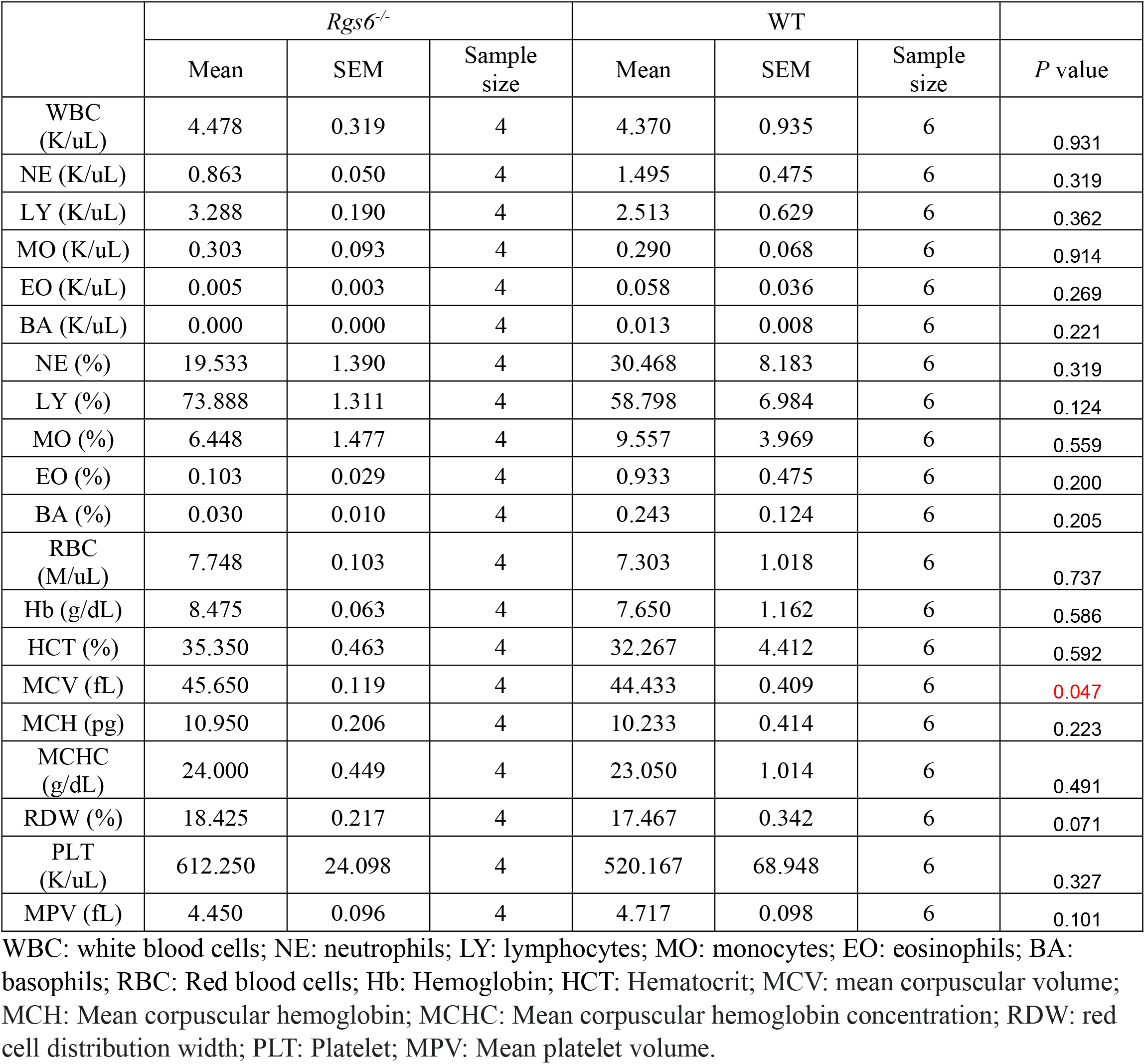
Whole blood cell counts in male WT and *Rgs6*^*-/-*^ mice.

|  | *Rgs6*<sup>-/-</sup> |  |  | WT |  |  | *P* value |
| --- | --- | --- | --- | --- | --- | --- | --- |
|  | Mean | SEM | Sample size | Mean | SEM | Sample size |  |
| WBC (K/uL) | 4.478 | 0.319 | 4 | 4.370 | 0.935 | 6 | 0.931 |
| NE (K/uL) | 0.863 | 0.050 | 4 | 1.495 | 0.475 | 6 | 0.319 |
| LY (K/uL) | 3.288 | 0.190 | 4 | 2.513 | 0.629 | 6 | 0.362 |
| MO (K/uL) | 0.303 | 0.093 | 4 | 0.290 | 0.068 | 6 | 0.914 |
| EO (K/uL) | 0.005 | 0.003 | 4 | 0.058 | 0.036 | 6 | 0.269 |
| BA (K/uL) | 0.000 | 0.000 | 4 | 0.013 | 0.008 | 6 | 0.221 |
| NE (%) | 19.533 | 1.390 | 4 | 30.468 | 8.183 | 6 | 0.319 |
| LY (%) | 73.888 | 1.311 | 4 | 58.798 | 6.984 | 6 | 0.124 |
| MO (%) | 6.448 | 1.477 | 4 | 9.557 | 3.969 | 6 | 0.559 |
| EO (%) | 0.103 | 0.029 | 4 | 0.933 | 0.475 | 6 | 0.200 |
| BA (%) | 0.030 | 0.010 | 4 | 0.243 | 0.124 | 6 | 0.205 |
| RBC (M/uL) | 7.748 | 0.103 | 4 | 7.303 | 1.018 | 6 | 0.737 |
| Hb (g/dL) | 8.475 | 0.063 | 4 | 7.650 | 1.162 | 6 | 0.586 |
| HCT (%) | 35.350 | 0.463 | 4 | 32.267 | 4.412 | 6 | 0.592 |
| MCV (fL) | 45.650 | 0.119 | 4 | 44.433 | 0.409 | 6 | 0.047 |
| MCH (pg) | 10.950 | 0.206 | 4 | 10.233 | 0.414 | 6 | 0.223 |
| MCHC (g/dL) | 24.000 | 0.449 | 4 | 23.050 | 1.014 | 6 | 0.491 |
| RDW (%) | 18.425 | 0.217 | 4 | 17.467 | 0.342 | 6 | 0.071 |
| PLT (K/uL) | 612.250 | 24.098 | 4 | 520.167 | 68.948 | 6 | 0.327 |
| MPV (fL) | 4.450 | 0.096 | 4 | 4.717 | 0.098 | 6 | 0.101 |

WBC: white blood cells; NE: neutrophils; LY: lymphocytes; MO: monocytes; EO: eosinophils; BA: basophils; RBC: Red blood cells; Hb: Hemoglobin; HCT: Hematocrit; MCV: mean corpuscular volume; MCH: Mean corpuscular hemoglobin; MCHC: Mean corpuscular hemoglobin concentration; RDW: red cell distribution width; PLT: Platelet; MPV: Mean platelet volume.

### RGS6 deficiency significantly enhanced ADP-induced platelet aggregation

We next investigated whether RGS6 affects platelet aggregation in response to two conventional platelet agonists, collagen and ADP. PRP was prepared from WT and *Rgs6*^*–/–*^ mouse blood by centrifugation, and platelet counts were normalized to 2.5 × 10^8^/mL using PPP. For each aggregation assay, 400 µL of the adjusted platelet suspension was used. As shown in **Fig. 2A**, collagen (1 µg/ml)-induced platelet aggregation was similar between WT and *Rgs6*^*–/–*^ platelets. In contrast, stimulation with ADP produced significantly greater platelet aggregation in *Rgs6*^*–/–*^ platelets than in WT controls (**Fig. 2B**). These findings suggest that loss of RGS6 preferentially augments ADP-mediated platelet activation rather than broadly increasing platelet responsiveness.

**Fig. 2.**
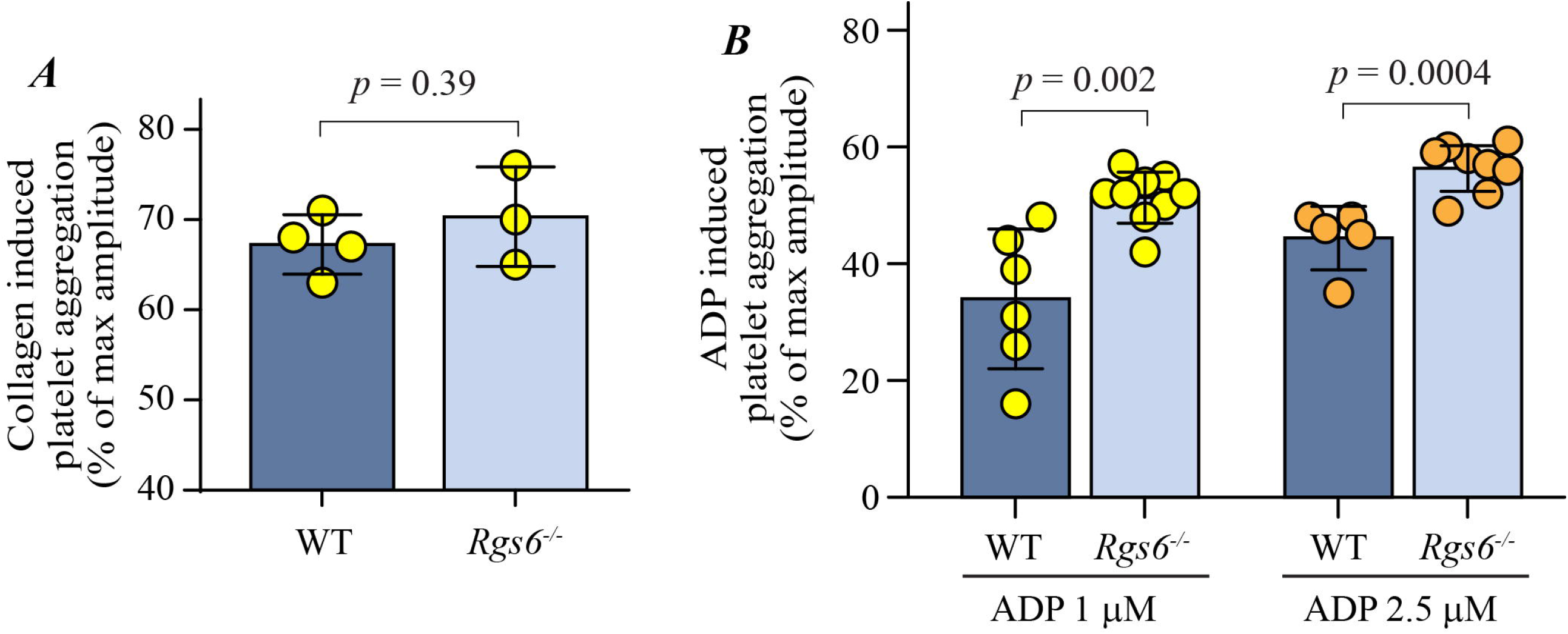
RGS6 deficiency enhances ADP-induced platelet aggregation in vitro. Whole blood was collected from WT and *Rgs6*^*–/–*^ mice using 0.109 M sodium citrate as an anticoagulant, Tyrode’s buffer (0.7 of blood volume) was added and PRP was prepared by centrifugation. Platelet concentration was adjusted to 2.5 × 10^8^ cells/mL with platelet-poor plasma (PPP). PRP (400 µL) was supplemented with CaCl_2_/MgCl_2_ to a final concentration of 1 mM, immediately before aggregation was initiated by addition of collagen (**A**) or ADP (**B**).

### RGS6 deficiency enhances ADP-induced suppression of the cAMP/PKA pathway in platelets

Since RGS6 is a potent negative regulator of Gi signaling [36] and P2Y_12_ is a well-defined Gi-coupled platelet GPCR [37], whereas coupling of PAR1 and PAR4 to Gi in platelets has not been clearly established [38], we hypothesized that RGS6 may serve as a key regulator of P2Y_12_ signaling. Consistent with this, ADP-induced platelet aggregation was significantly enhanced in *Rgs6*^*–/–*^ platelets (Fig. 2B). To further investigate this, we examined the effect of RGS6 on platelet signaling in response to ADP stimulation. Platelets pooled from 6 WT and 6 *Rgs6*^*–/–*^ mice were aliquoted and stimulated with ADP over a time course and effectors of the P2Y_12_/Gi signaling axis were analyzed by Western blot.

As shown in **Fig. 3A**, RGS6 deficiency did not affect P2Y_12_ expression on platelets. Phosphorylation of AKT, a downstream effector of PI3K and a common marker of P2Y_12_-dependent signaling [31, 39], increased rapidly in WT platelets but was delayed and remained at low levels in *Rgs6*^*–/–*^ platelets (Fig. 3A & **3B**). In contrast, phosphorylation of PKA (**Fig 3C & 3D**) and its substrate vasodilator□stimulated phosphoprotein phosphorylation (p-VASP, Fig. 3C and **3E**), a well-established readout of cAMP/PKA signaling and an inverse marker of P2Y_12_ activity [40], were markedly reduced in *Rgs6*^*–/–*^ platelets following ADP stimulation, indicating enhanced suppression of the cAMP/PKA pathway. Together, these data indicate that RGS6 deficiency alters the profile of ADP-induced P2Y_12_ signaling. Specifically, loss of RGS6 enhances Gi-mediated suppression of the inhibitory cAMP/PKA pathway.

**Fig. 3.**
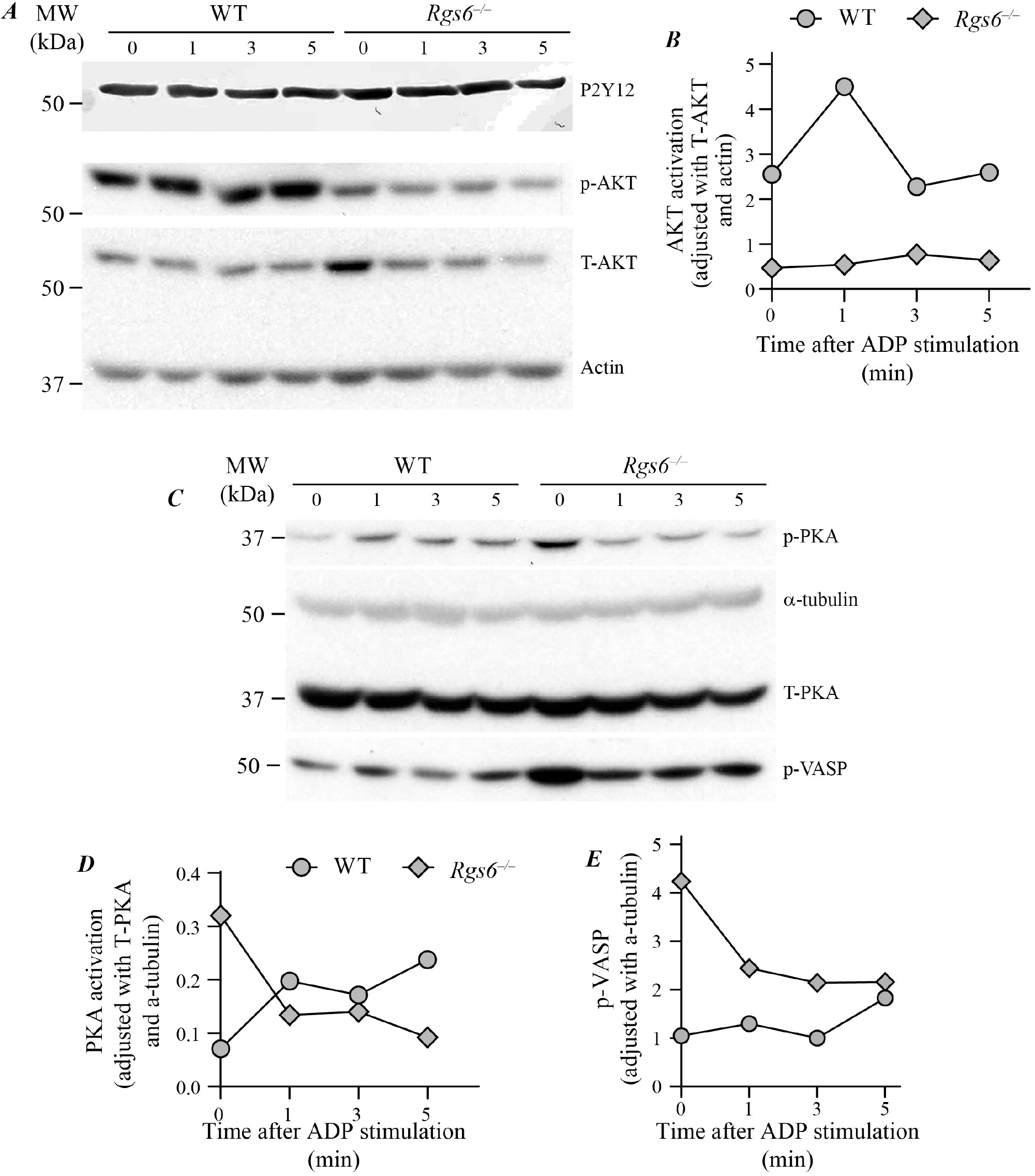
RGS6 deficiency enhances ADP-mediated suppression of cAMP/PKA pathway. Platelet-rich plasma (PRP) pooled from 6 mice per genotype was divided into 4 aliquots and stimulated with 2.5 μM ADP for 0, 1, 3, and 5 min. Reactions were terminated by addition of EDTA (1 mM final) and PGEL (0.5 μM final). Platelets were immediately pelleted by centrifugation and lysed in RIPA buffer containing protease and phosphatase inhibitors. Equal amounts of protein (30 μg) were subjected to Western blot analysis using the indicated antibodies. Band intensities were quantified using ImageJ. Data are presented as the ratio of phosphorylated (p-) protein to total (T-) protein, further corrected for loading by dividing by the relative loading-control intensity normalized to WT at time 0. **A&B**: GβγPI3K/AKT signaling, **C, D, and E:** Gai/AC/cAMP/PKA signaling.

### RGS6 modulates P2Y_12_/G_i_ signaling

To further demonstrate that RGS6 regulates platelet ADP/P2Y_12_ signaling, we performed a cAMP reporter assay using HEK293 cells expressing a luciferase reporter under the control of a cAMP-responsive element (CRE). Plasmids encoding murine RGS6 and P2Y_12_ (and Renilla luciferase) were transfected into HEK293 cells and luciferase/Renilla activity was measured following ADP stimulation. Data from all experimental groups were normalized to cells transfected with empty vector (pcDNA3.1) alone.

Forskolin treatment induced more than a 10 fold increase in luciferase activity (**Supplemental Figure II**), confirming that the assay system was functional. Although HEK293 cells have been reported to express P2Y receptors [41], ADP did not significantly affect luciferase activity in cells transfected with pCDNA3.1 (**Fig. 4**, blue bar, *p* = 0.28), suggesting limited functional endogenous P2Y signaling under these conditions. Overexpression of P2Y_12_ resulted in a modest reduction in luciferase activity following ADP stimulation (light blue, *p* = 0.089), consistent with Gi-mediated inhibition of cAMP signaling. Expression of RGS6 alone significantly increased basal level luciferase activity, which is reduced by ADP treatment (brown bar, *p* = 0.033), supporting the presence of low-level endogenous Gi signaling. Co-expression of RGS6 with P2Y_12_ significantly increased luciferase activity compared with P2Y_12_ alone at all ADP concentrations tested (0, 10, and 20 μM; *p* = 0.0009, 0.008, and 0.02, respectively), indicating attenuation of P2Y_12_ signaling by RGS6.

**Fig. 4.**
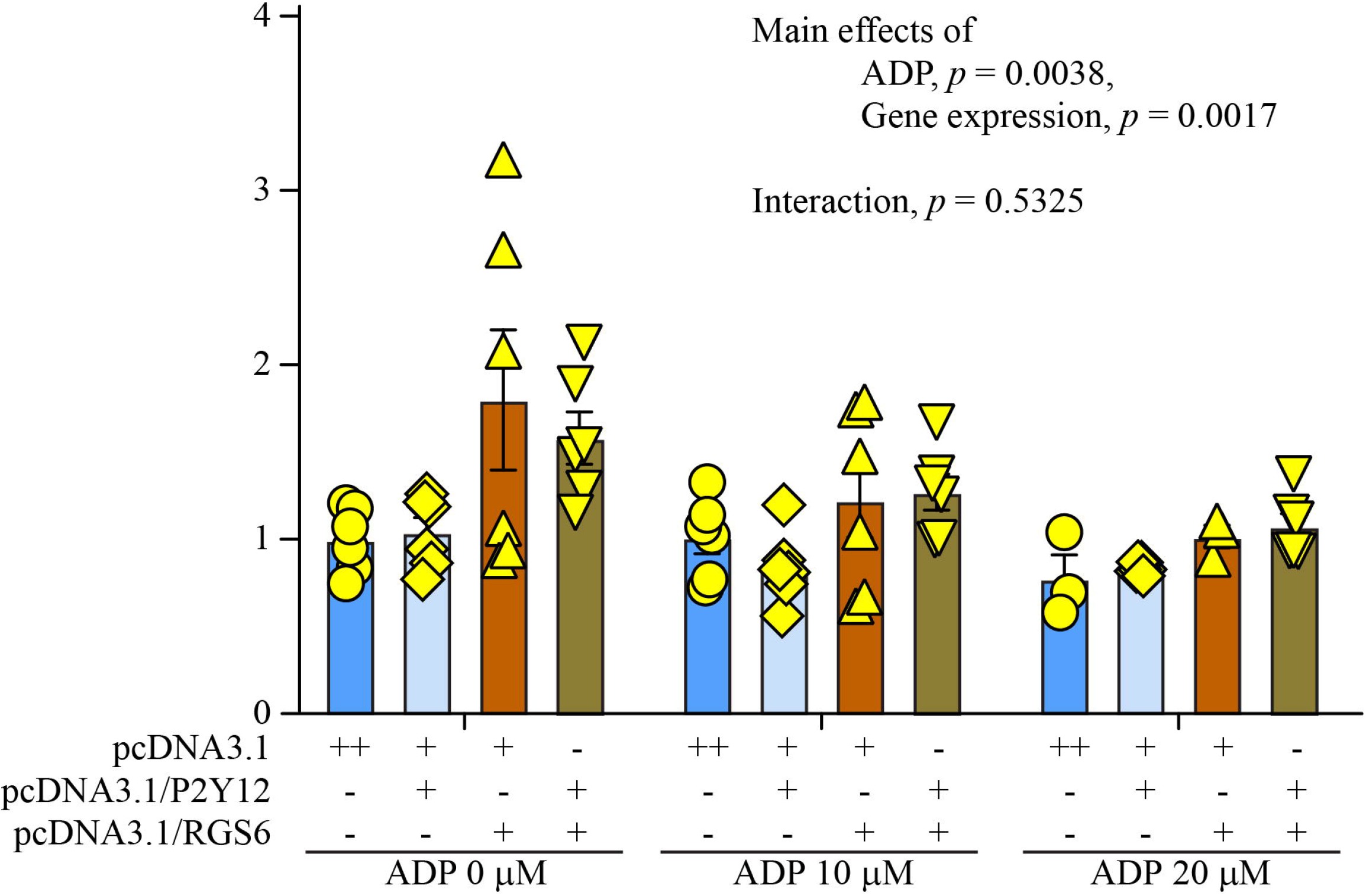
RGS6 negatively regulates P2Y12/Gi signaling. GloResponse™ CRE-luc2P HEK293 cells were seeded in 24-well plates at a density of 1 × 10^2^ cells per well. Cells were transfected with the indicated plasmids 6 h after seeding and incubated for an additional 24 h. Cells were then serum-starved for 4 h prior to stimulation with ADP at the indicated concentrations. “+” indicates 0.25 µg plasmid.

Two-way ANOVA demonstrated significant main effects of ADP (*p* = 0.0038) and gene expression (*p* = 0.0017), with no significant interaction (*p* = 0.533), suggesting that the effect of RGS6 on P2Y_12_ signaling is consistent across agonist concentrations. Together, these data support a role for RGS6 as a negative regulator of P2Y_12_-dependent Gi signaling.

### RGS6 regulates in vivo thrombosis through a P2Y_12_-dependent mechanism

To determine whether the prothrombotic phenotype observed in *Rgs6*^*–/–*^ mice was functionally dependent on P2Y_12_/G_i_ signaling, WT and *Rgs6*^*–/–*^ mice were treated with clopidogrel (1 mg/kg) once per day for 72h and then subjected to the FeC□ -induced carotid artery injury model [31]. As shown in **Fig. 5**, clopidogrel treatment markedly prolonged the time to form an occlusive thrombus in both strains and diminished the difference in thrombotic responses between male WT and *Rgs6*^*–/–*^ mice. These findings demonstrate that the prothrombotic phenotype associated with RGS6 deficiency is largely dependent on P2Y_12_ signaling, consistent with the role of RGS6 as a regulator of Gi. These findings support the interpretation that the prothrombotic phenotype associated with RGS6 deficiency is largely P2Y_12_-dependent.

**Fig. 5.**
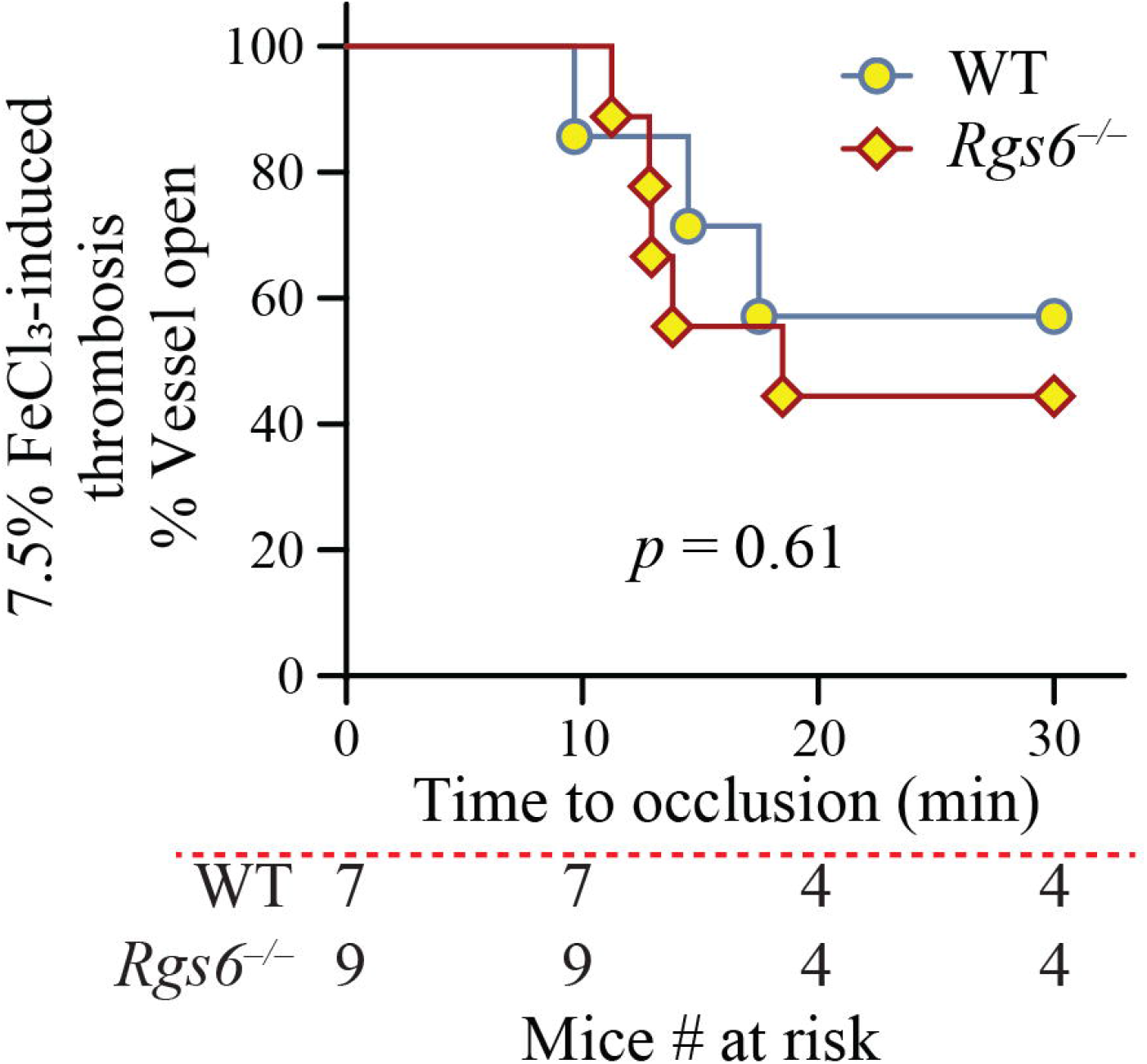
Inhibition of P2Y_12_ with clopidogrel abolishes the enhanced arterial thrombosis observed in *Rgs6*^*–/–*^ mice. WT and *Rgs6*^*–/–*^ mice were administered clopidogrel (1 mg/kg/day) by oral gavage for 3 days prior to induction of thrombosis.

## Discussion

RGS proteins regulate GPCR by accelerating the rate of GTP hydrolysis by Gα subunits [42]. RGS6 has not been extensively studied in platelets. In this study, we identified RGS6 as a negative regulator of platelet-intrinsic arterial thrombosis. Genetic deletion of *Rgs6* accelerated thrombus formation in vivo and enhanced ADP-induced platelet aggregation ex vivo. Importantly, pharmacologic inhibition of P2Y_12_ with clopidogrel abolished the genotype-dependent difference in thrombosis in *Rgs6*^*–/–*^ and WT mice, indicating that the prothrombotic phenotype associated with RGS6 deficiency is P2Y_12_-dependent.

Our findings are consistent with clinical observations in COVID-19, a condition associated with increased thrombotic risk and reduced platelet RGS6 expression (∼30%) [23]. In addition, a prior study identified the rs9323567C>T SNP within an intronic region of RGS6.

Although RGS6 expression was not examined in that study, such variants are often associated with reduced gene function. Our finding may therefore provide a mechanistic explanation for the reported association between the RGS6 rs9323567C>T variant and decreased P2Y_12_ Reaction Units (PRU) in patients treated with clopidogrel [27]. Moreover, these results raise the possibility that reduced RGS6 activity may contribute to variability in P2Y_12_ inhibitor responsiveness, which could partially explain the limited clinical benefit observed in studies of COVID-19-related mortality [43].

Importantly, the cAMP-responsive luciferase assays demonstrated an inverse, dose-dependent relationship between ADP and P2Y_12_/Gi signal, increased luciferase activity in the presence of RGS6, and no significant interaction between RGS6 expression and the dose-response. These data support our in vitro signaling findings, indicating that RGS6 suppresses the P2Y_12_/Gi pathway. The lack of a significant RGS6 x ADP interaction suggests that RGS6 does not alter agonist sensitivity but instead modulates the overall magnitude of P2Y_12_/Gi signaling. This interpretation is consistent with our in vivo thrombosis studies: under basal conditions, loss of RGS6 increases the ADP/P2Y_12_ signal output, thereby accelerating occlusive thrombus formation (**Fig. 6**), whereas pharmacologic P2Y_12_ blockade using clopidogrel abolishes this difference.

**Fig. 6.**
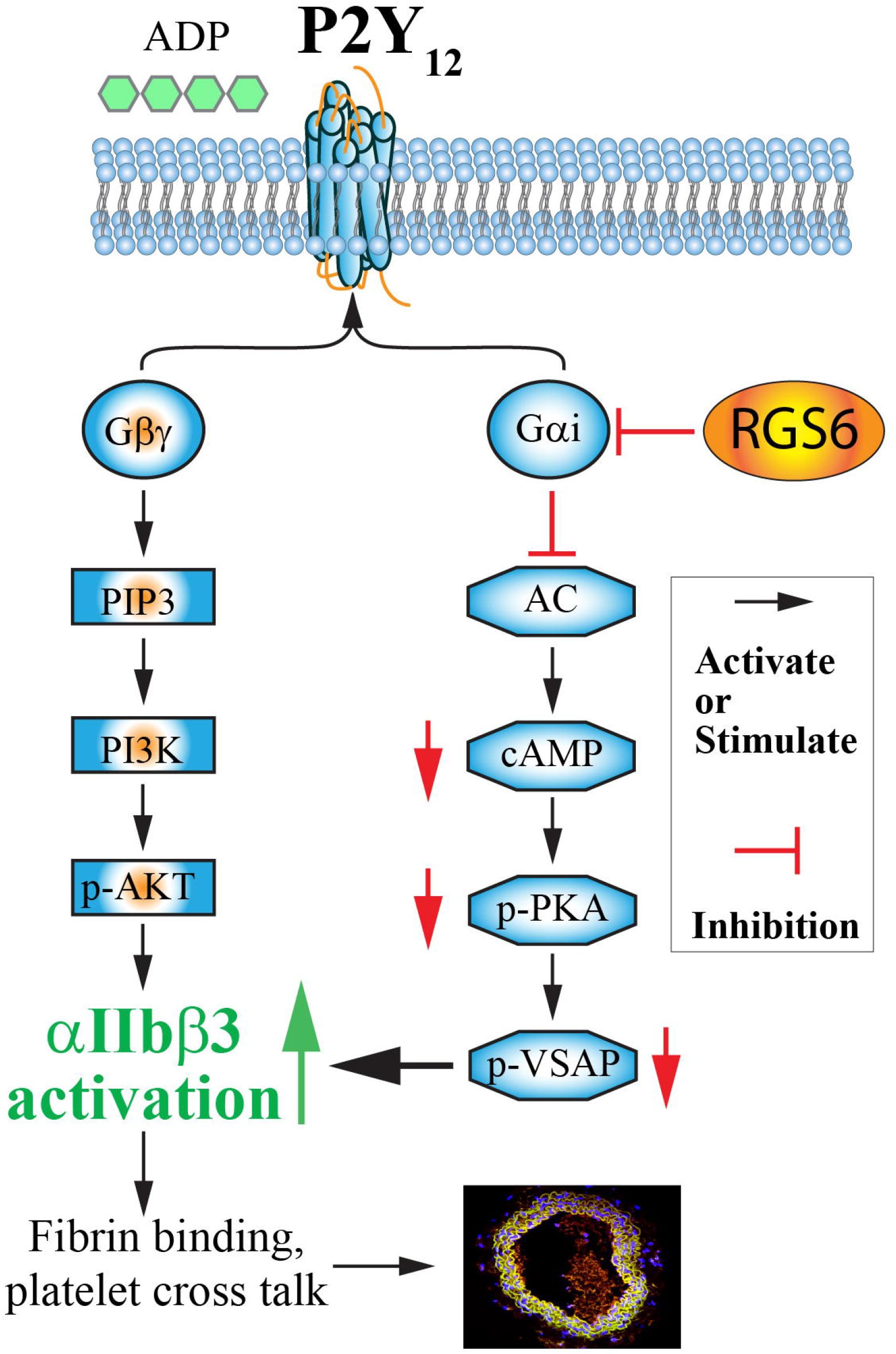
Schematic of RGS6-mediated negative regulation of P2Y12/Gi signaling. Black arrows indicate activation, and blue blunt-ended lines indicate inhibition. AC, adenylyl cyclase.

In addition to RGS6, other RGS proteins, including RGS10 [44], RGS18 [19], and RGS16 [21, 45], have been reported to target Gi and regulate platelet activation (**Table 2**). Although ADP-induced aggregation was increased in *Rgs6*^*–/–*^ platelets, AKT phosphorylation was delayed and reduced. However, PKA and VASP phosphorylation were reduced at all time points following ADP stimulation. These findings suggest that RGS6-mediated modulation of the P2Y_12_/Gi/cAMP/PKA pathway may play a more prominent role than Akt signaling in regulating platelet activation.

**Table 2:** Summary of RGS proteins with evidence on platelet activation.

| **RGS protein** | **Gi activity** | **Platelet evidence** | **Deficiency on Akt (PI3K/Gβγ)** | **Deficiency on PKA / VASP (cAMP)** | **Interpretation** |
| --- | --- | --- | --- | --- | --- |
| **RGS10** | Strong | Strong | ↓ Akt | ↑ PKA / ↑ VASP | Classic Gi inhibition → less PI3K, more cAMP |
| **RGS18** | Moderate | Strong | ↓ Akt (context-dependent) | ↑ PKA / ↑ VASP | Broad regulator; modulates both Gi and Gq, net inhibitory |
| **RGS6** | Strong (biochemical) | Emerging | ~ no consistent ↑ (context-dependent; may ↓ or unchanged) | ↑ PKA / ↑ VASP (baseline effect) | Kinetic regulator of Gi; strongest effect on cAMP/PKA, variable effect on Akt |
| **RGS16** |  | limited | Likely ↓ Akt (limited data) | Likely ↑ PKA | Expected Gi GAP behavior, but not well defined in platelets |

Our results also support a potential sex-dependent role for RGS6 in thrombosis. Female *Rgs6*^*–/–*^ mice exhibited a trend opposite to that observed in males, with prolonged occlusion times compared with WT mice. In our recent study, we found that P2Y_12_ forms complexes with the Na^+^/K^+^-ATPase α1 subunit, and α1 haplodeficiency significantly inhibits thrombosis in males but not in females [32]. Given that female platelets express higher levels of α1 than male platelets, and female platelet are more sensitive to ADP [43], P2Y_12_ signaling may be regulated in a context-dependent manner. These findings raise the possibility that RGS6 contributes to sex-specific regulation of platelet function, which warrants further investigation.

Members of the RGS family exhibit differences in their tissue distribution and have different specificities for various Gα isoforms [16]. Consequently, RGS proteins have emerged as potential new therapeutic targets. RGS inhibitors may provide a way to enhance the effects of endogenous agonists or may be combined with exogenous agonists to produce responses with greater tissue selectivity than can be achieved with an agonist alone [46]. Prior work suggests that RGS6 may provide a therapeutic target for the treatment of alcohol use disorder [47, 48], Parkinson’s disease [49], and cardiac ischemic injury [50]. However, the data presented here suggest that pharmacotherapies that suppress RGS6 function may be limited by an increased risk of thrombosis-related morbidities such as myocardial infarction or ischemic stroke.

This study has several limitations. First, the lack of a reliable antibody for RGS6 limited direct assessment of protein expression, and mechanistic conclusions relied on downstream signaling readouts and a heterologous reporter system. While informative, this system represents a reductionist model and does not fully recapitulate the complexity of platelet signaling. Second, *Rgs6* deletion was global, and contributions from non-platelet cell types cannot be excluded.

However, the enhanced ADP-induced aggregation observed ex vivo and the normalization of thrombosis by P2Y_12_ inhibition strongly support a platelet-intrinsic mechanism.

## Conclusion

We identify RGS6 as a previously unrecognized negative regulator of platelet-intrinsic arterial thrombosis. Loss of RGS6 accelerated thrombus formation in vivo, enhanced ADP-induced platelet aggregation ex vivo, and altered signaling within the platelet ADP/P2Y_12_ axis. cAMP reporter assays and pharmacologic inhibition studies indicate that RGS6 modulates the magnitude of P2Y_12_/Gi signaling rather than altering agonist sensitivity. Together, these findings establish RGS6 as an important regulator of platelet function and thrombosis.

## Supporting information

none

## Author’s contributions

Conception and design: WL, BR

Data acquisition and analysis: RR, BR, WL, RJ, HY, SC Data interpretation: RR, BR, WL.

Manuscript drafting: RR, WL.

Critical revision of the manuscript: RR, BR, HY, SC. Final approval of the version to be published: WL.

## Acknowledgements

We thank Dr. Rory Fisher (University of Iowa School of Medicine) for providing the *Rgs6*^*–/–*^ mice used in this study.

