## Supplementary material for "Regulator of G Protein Signaling 6 Negatively Regulates Platelet Activation and Arterial Thrombosis in Mice": none

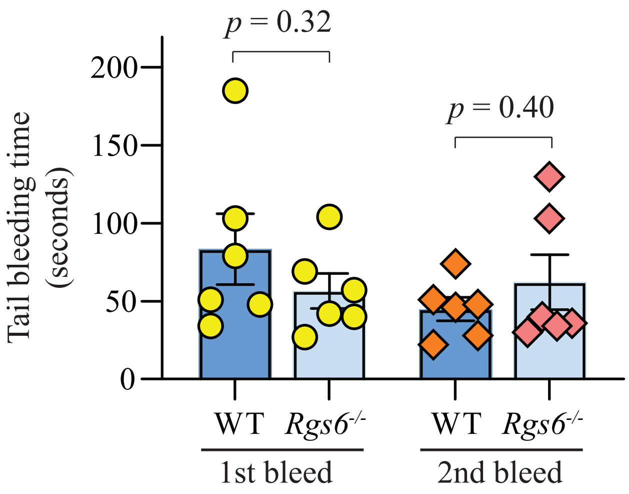


**Supplemental Figure I. RGS6 deficiency does not affect bleeding time in mice.** Tail bleeding assays were performed in anesthetized mice by transecting 1 cm from the tail tip, and bleeding time was recorded until cessation of bleeding.

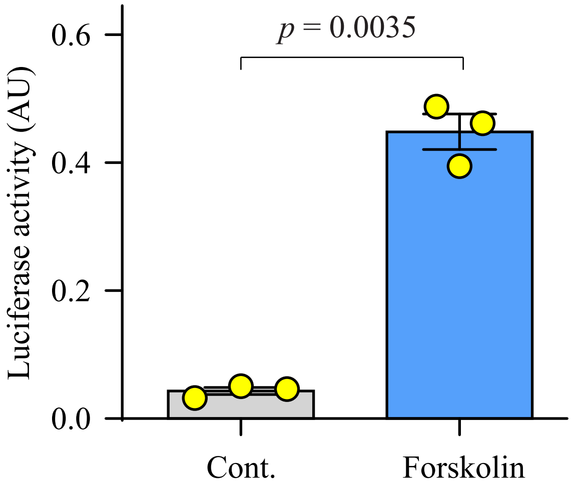


**Supplemental Figure II. Forskolin treatment significantly increased luciferase activity in HEK293 cAMP reporter cells.**
